# Simulated Microgravity Recapitulates Aspects of Biological Aging in Humans

**DOI:** 10.64898/2026.03.03.709404

**Authors:** Fei Wu, Alexander Chebykin, Anna Rychkova, Kevin Schneider, Matias Fuentealba, Abhayjit Saini, Heather Halaweh, Nathan Bracey, Mark Davis, Daniel A. Winer, David Furman

**Author notes:** Corresponding author David Furman, PhD, Professor, Buck Institute for Research on Aging.

## Abstract

Spaceflight and microgravity profoundly affect human physiology and have been proposed to recapitulate key features of biological aging, yet the underlying mechanisms remain incompletely understood. Here, we performed whole-genome transcriptomic profiling to define immune cell alterations associated with both natural aging and simulated microgravity. Leveraging the longitudinal nature of the Stanford 1,000 immunomes Project, we compared peripheral blood mononuclear cells (PBMCs) exposed to rotating wall vessel bioreactor with matched samples collected up to 9 years later from the same individuals. We quantified changes across aging hallmarks, molecular pathways, gene modules, cellular energetics, disease risk and vaccine-response signatures. Microgravity-induced transcriptional closely tracked subject-level aging trajectories spanning across disease risk domains including those affecting the metabolic, musculoskeletal and circulatory systems, and multiple aging hallmarks involving nutrient sensing, intrinsic capacity, chronic inflammation, proteostasis, cellular senescence and metabolic regulation. Independent validation using Single-Cell Energetic Metabolism by Profiling Translation Inhibition (SCENITH) profiling confirmed these observed metabolic adaptations and revealed reduced mitochondrial dependence with minimal compensatory glucose dependence across immune cell subsets, features that strongly parallel aging biology. Consistent with previous findings, longitudinal changes indicated that close of 1/3 of participants do not follow population trajectories but these can be partly predicted with simulated microgravity exposure. Together, this within-donor framework establishes simulated microgravity as a scalable and experimentally tractable platform to model aspects of biological aging in humans and accelerating the prioritization of candidate countermeasures for spaceflight and aging on Earth.

## Introduction

Spaceflight and microgravity act as system-wide stressors, perturbing immune, musculoskeletal, cardiovascular and metabolic physiology with implications for astronaut health^1–3^. Interpreting these multifactorial responses requires integrative approaches spanning biological scales. Because single pathways rarely capture whole-organism adaptation, approaches that connect system-level phenotypes to coordinated pathway programs and gene-level regulation are warranted. We and others have shown that biological aging is affected during spaceflight, and these changes can be emulated on Earth by exposing cells and tissues to ground analogs of microgravity ^4,5^, motivating the use of aging as a comparative reference for interpreting microgravity-associated responses ^6–9^. This evidence supports a multi-scale view that connects systems-level phenotypes to molecular and cellular energetics. Such an approach ensures that conclusions rely on consistent signal alignment across biological scales rather than isolated biomarkers.

Prior studies in spaceflight, ground-based analogs, and aging report overlapping signals, including mitochondrial stress, immune remodeling and mechanotransduction/cytoskeletal programs ^4,10–12^. However, these rely on cross-sectional data and the use of the so-called “biological aging clocks” as surrogate endpoints which capture a partial picture (population trends) and not subject-level biological trajectories. Direct within-donor comparisons that quantify microgravity-induced shifts alongside aging-associated changes, integrate systems-to-pathways-to-genes, and anchor transcriptomic concordance with functional peripheral blood mononuclear cell (PBMC) metabolism remain uncommon ^10,13^. Consequently, it remains unclear which biological programs show coherent donor-level coupling across scales and whether transcriptomic concordance aligns with functional energetic phenotypes. This uncertainty complicates efforts to distinguish broadly shared stress responses from context-specific adaptations that could require distinct countermeasures.

Here we apply a fully paired, within-donor design using a longitudinal immune aging cohort (Stanford 1,000 Immunomes Project) to test whether simulated microgravity-induced shifts covary with longitudinal aging-associated changes across system-level categories and molecular programs. We assessed whether an orthogonal Peripheral Blood Mononuclear Cells (PBMC) energetic assay could provide compatible functional context using Single-Cell Energetic Metabolism by Profiling Translation Inhibition (SCENITH) profiling^14–17^. Our results indicate that exposure of PBMC to simulated microgravity emulates multiple components of natural aging observed in the longitudinal cohort including disease risk biomarkers, aging pathways, and vaccine response profiles.

## Results

### Cohort-level coupling across aging hallmark signatures and DiseaseAge domains

To quantify the alignment between spaceflight-like stress and biological aging, we computed paired, within-donor delta changes relative to a shared baseline. We defined ΔμG as the within-donor shift induced by 24h simulated microgravity (Baseline μG – Baseline 1G) (as described previously^4^), and ΔAging as the longitudinal aging trajectory (Natural aging 1G – Baseline 1G) (Fig. 1A). We tested the concordance of delta change first at the system level using aging hallmark gene sets (see Methods) and DiseaseAge^18^ domains and subsequently at molecular and functional levels in downstream analyses.

**Fig. 1.**
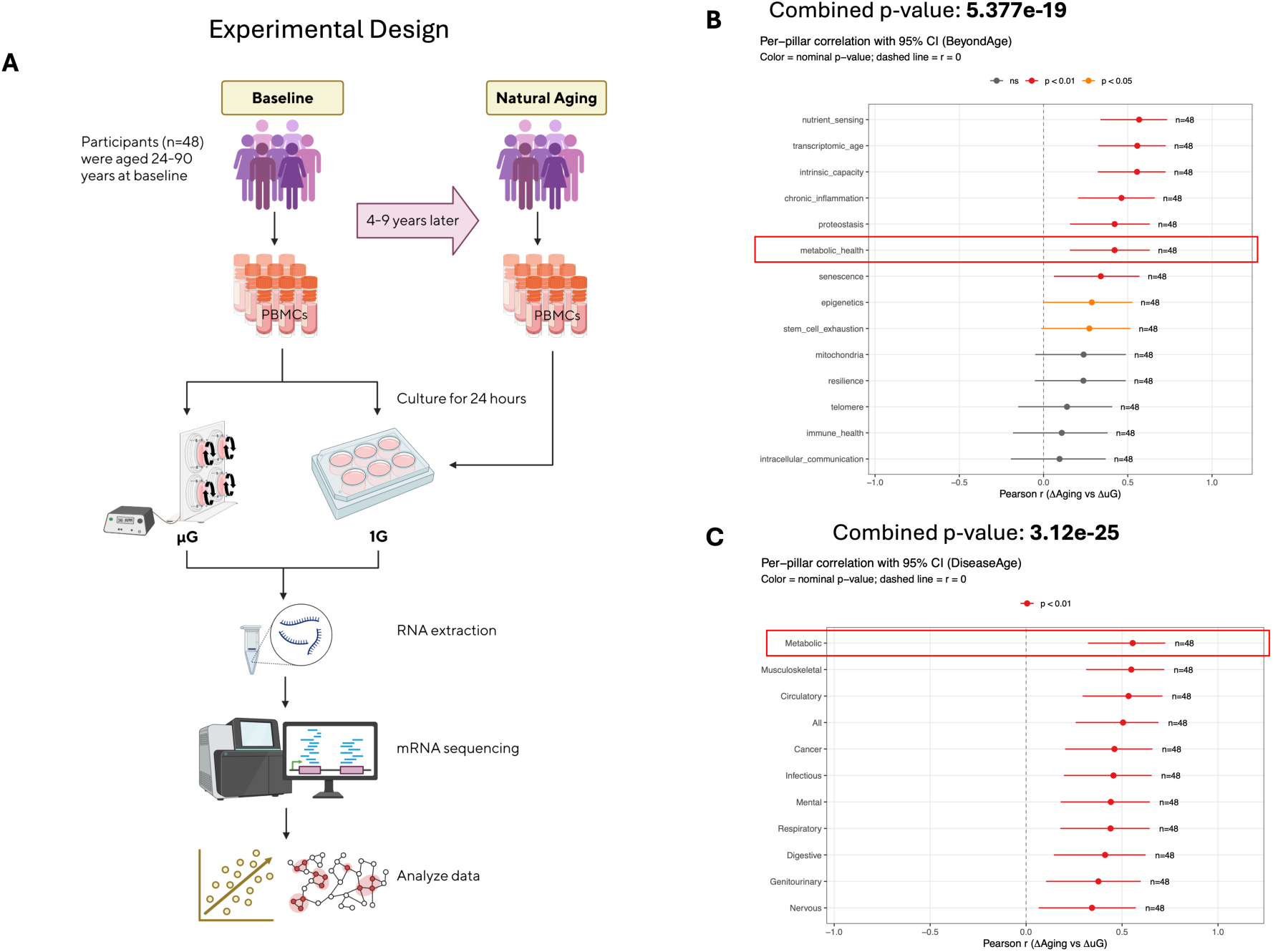
Cohort-level coupling between simulated microgravity and longitudinal aging in a paired human design. (A) Study design schematic for the 1KIP cohort (n=48). PBMCs were profiled at Baseline (1G), after simulated microgravity exposure (μG), and at longitudinal follow-up under 1G (“Aging”). Within each donor, deltas were computed relative to Baseline: ΔμG = (μG − Baseline) and ΔAging = (Aging − Baseline). (B) BeyondAge pillar correlation. Forest plot summarizing Pearson correlation coefficients (r) between donor-level ΔμG and ΔAging for each BeyondAge pillar. Points represent r values with 95% confidence intervals; colors indicate nominal significance thresholds (p < 0.05, p < 0.01). The dashed vertical line marks r = 0. A combined p-value across all pillars was computed using Fisher’s method. (C) DiseaseAge domain correlation. Analogous forest plot for DiseaseAge domains, showing correlations between donor-level ΔμG and ΔAging with 95% confidence intervals and Fisher’s combined p-value. Donor-level Δ values underlying panels (B–C) are visualized in Supplementary Fig. 1 and tabulated in Supplementary Tables 1–2.

To test whether simulated microgravity and longitudinal aging induce aligned system-level shifts, we estimated 14 transcriptomic aging hallmarks derived from independent models and anchored to the GTEx^19^ population reference (n=803). Across the 48-donor from Stanford 1,000 immunomes Project (1KIP) paired cohort, ΔμG was positively coupled with ΔAging across multiple aging hallmarks, consistent with broad concordance at the system level (Fig. 1B; donor-level scatter plots in Supplementary Fig. 1A). Nine of 14 pillars were nominally significant, led by nutrient_sensing (r=0.567, p=1.30e-5), transcriptomic_age (r=0.556, p=2.01e-5), and intrinsic_capacity^20^(r=0.555, p=2.13e-5), with chronic_inflammation also showing a robust association (r=0.463, p=4.65e-4). Notably, the metabolic_health pillar was also positively correlated (r=0.422, p=0.00142). The composite summary (’All’) was positive (r=0.465, p=4.40e-4), and the combined p-value across the 14 pillars was highly significant (combined p=5.38e-19; Fisher’s method). Together, these results confirm a robust concordance between microgravity-induced shifts and aging trajectories.

These results prompted us to determine whether this alignment extends to specific clinical risks signatures. To do so, we relied on the DiseaseAge models corresponding to transcriptomic signatures of mortality risk estimators mapped to ICD-10 chapters, recently reported by our group^18^.

Analysis of DiseaseAge domains^18^ confirmed this extension, where all 11 categories reached nominal significance (Fig. 1C; Supplementary Fig. 1C). The strongest associations emerged in the Metabolic (r=0.555, p=2.09e-5) and Musculoskeletal (r=0.548, p=2.77e-5) domains, followed closely by Circulatory system (r=0.534, p=4.64e-5). The composite summary (‘All’) retained this positive signal (r=0.505, p=1.25e-4), yielding a robust combined significance across all domains (combined p=3.12e-25, Fisher’s method). Detailed donor-level values for these correlations are provided in Supplementary Table 1-2.

To quantify how much donor-to-donor variation in ΔAging is captured by ΔμG at the systems level, we performed a complementary predictive analysis using univariate regressions evaluated by leave-one-out cross-validation. Using the aging hallmark signatures “All” and DiseaseAge “All” aggregates, ΔμG predicted ΔAging with cross-validated accuracies of 84% and 87%, respectively (Supplementary Fig. 1B,D; donor-level scatterplots in Supplementary Fig. 1A,C). These results confirm that the positive correlation between microgravity-induced shifts and aging trajectories spans diverse disease-linked physiological systems and can partially predict aging trajectories. Having established this robust system-level concordance, we next examined whether this coupling extends to molecular resolution using pathway-level analysis.

### Global pathway concordance between ΔAging and ΔμG shifts

To assess concordance at the molecular system level, we aggregated 14,500 gene sets spanning three major MSigDB pathway collections: (1) Hallmark, (2) Curated Canonical Pathways (C2:CP), and (3) Gene Ontology (C5:GO). Across this combined collection, single-sample gene set enrichment analysis (ssGSEA)^21^ revealed a global positive shift in correlations between ΔAging and ΔμG, indicating widespread concordance across donors (Fig. 2A). The distribution centered at a mean correlation of 0.354 (median = 0.360) with an interquartile range (IQR) of 0.181. Notably, 99.3% of pathways exhibited positive coupling, and 66.9% exceeded a correlation threshold of 0.3. This global shift suggests that the alignment between microgravity and aging is not limited to specific aging pathways but reflects broader similarities across the transcriptome.

**Fig. 2.**
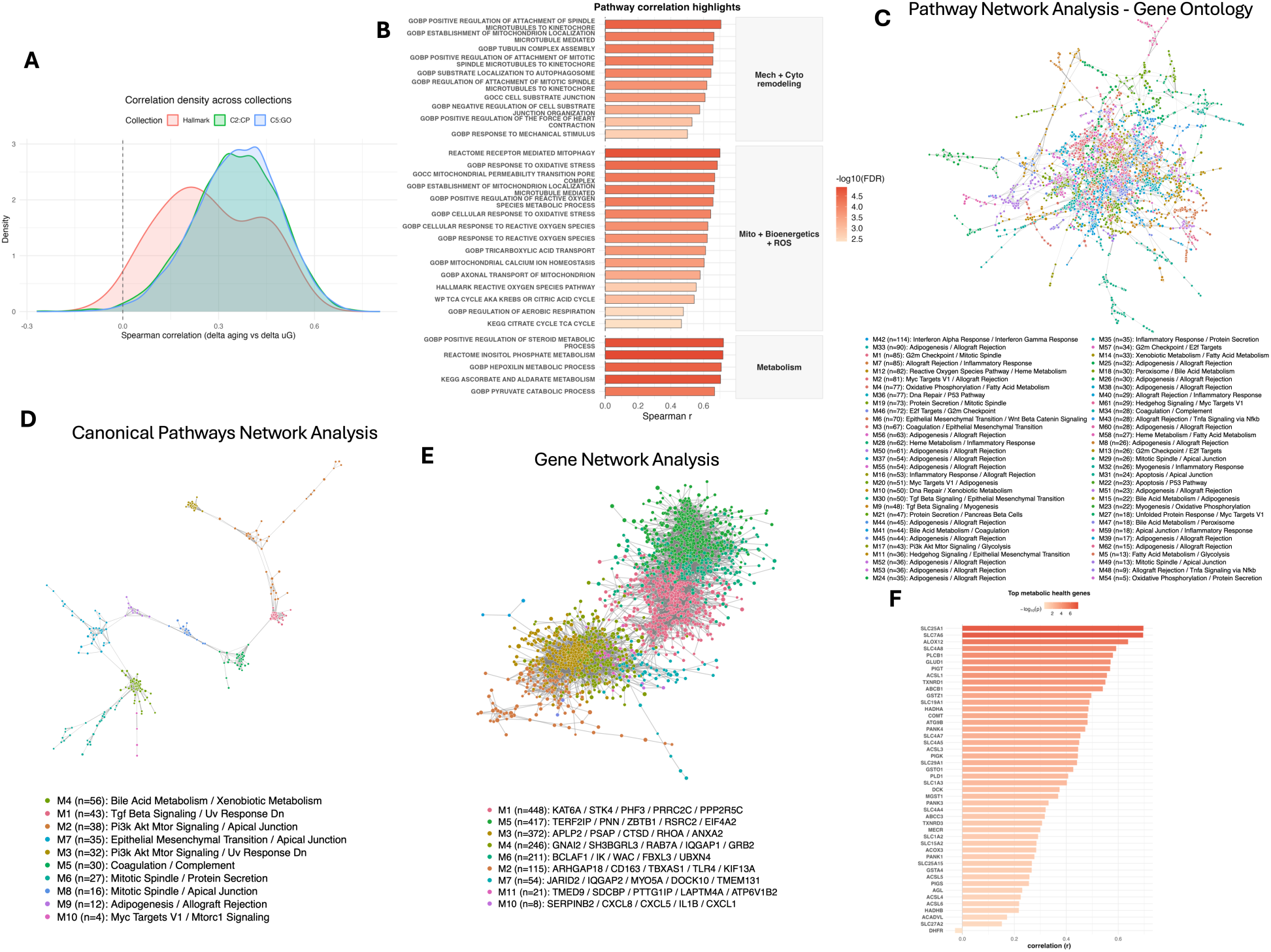
Global pathway concordance and network organization of coupled aging and simulated microgravity programs. (A) Density distributions of pathway-wise Spearman correlations between ssGSEA delta scores (ΔAging vs Baseline and ΔμG vs Baseline) across Hallmark (n=50), C2 (n=4,014), and C5 (n=10,436) collections. Curves show the distribution of correlations for each collection; the dashed vertical line marks zero. (B) Theme-level aggregation of pathway coupling. Bar plot shows representative pathways from mechanotransduction, cytoskeleton, mitochondrial organization/bioenergetics, ROS, and broad metabolism, selected as the top 5 per theme by Spearman ρ among significant pathways. Bar height is Spearman’s correlation coefficient between ΔAging and ΔμG across 48 donors; fill encodes −log10(FDR). Full pathway lists and statistics are provided in Supplementary Tables 3–7. (C) Pathway network for C5. Nodes are pathways with significant correlation (Spearman) of ΔAging and ΔμG, sized by |ρ|. Edges are weighted by Jaccard gene-set overlap and visualized after trimming to the top 25% of edge weights. Colors denote Louvain modules within the largest connected component. Module labels reflect the top two Hallmark pathways per module from weighted Hallmark enrichment. (D) Pathway network for C2, plotted and annotated as in panel (C). (E) Gene-level network modules. Genes with significant across-donor coupling between ΔμG and ΔAging were used to build a similarity network based on combined delta profiles; Louvain community detection identified modules that were annotated by Hallmark enrichment and module-level coupling of ΔμG and ΔAging. (F) Metabolic-health pillar gene anchors. Within the BeyondAge metabolic-health pillar gene set, per-gene ΔμG and ΔAging values were correlated across donors. Highlighted genes illustrate concordant nutrient transport and mitochondrial/lipid signaling anchors.

This positive shift was robust across the large gene ontology collections. The Gene Ontology library (n=10,436) displayed a mean correlation of 0.356 (99.4% positive), while the Curated Canonical Pathways library (n=4,014) showed a mean of 0.350 (99.1% positive). The smaller Hallmark collection (n=50) also skewed positive (98% of pathways) but displayed more modest effect sizes (mean correlation = 0.272; 38% > 0.3). This suggests that concordance is robust in broad across ontology and curated pathway collections. Full correlation statistics for all pathways are provided in Supplementary Tables 3–6. These general collection-level distributional shifts establish the rationale for a more detailed investigation of specific biological themes and network modules.

### Theme-level concordance and pathway network modules organize coupling between ΔμG and ΔAging

To gain insights into specific mechanisms involved, we organized the global pathway signal by aggregating pathways into broad functional themes. This revealed robust concordance across mechanotransduction, cytoskeleton, mitochondria, reactive oxygen species (ROS), and metabolism (Fig. 2B, Supplementary Table 7). Structural themes were highly consistent: mechanotransduction included 36 significant pathways (all positive; median correlation = 0.401), while cytoskeleton comprised 232 positive pathways (median correlation = 0.415). Mitochondrial and oxidative stress programs showed similar alignment, with 115 mitochondrial and 69 ROS pathways all exhibiting positive correlations. Notable examples driving this signal include GOBP_SUBSTRATE_LOCALIZATION_TO_AUTOPHAGOSOME (r = 0.645) and REACTOME_RECEPTOR_MEDIATED_MITOPHAGY (r = 0.700), highlighting a link between structural remodeling and mitochondrial quality control during subject level aging and simulated microgravity exposure. This axis is further reinforced by the robust alignment of GOBP_RESPONSE_TO_OXIDATIVE_STRESS (r = 0.684), identifying oxidative pressure as a key component of this remodeling. Metabolic themes were also widely represented, including 17 specific energy/bioenergetics pathways (e.g., GOBP_TRICARBOXYLIC_ACID_TRANSPORT, r = 0.612) and over 800 broad metabolic terms.

Cross-referencing these specific molecular pathways against the Hallmark collection supports a specific, stress-linked interpretation of this metabolic signal (Fig. 2B). At the Hallmark level, while the Reactive Oxygen Species Pathway remained significant (r = 0.555, FDR = 0.002), the broad Oxidative Phosphorylation and Glycolysis sets were null or modest (r = 0.030 and r = 0.291, respectively). This pattern suggests that metabolic coupling is significantly affected regardless of the pathway level analysis, it is distributed across multiple granular Gene Ontology and Curated Pathway analysis, such as those governing ROS handling and mitochondrial maintenance.

Additional clustering analysis showed these concordant pathways can be assembled into coherent functional modules (Fig. 2C–D, Supplementary Table 8-9) with the C5 network clustering into Hallmark-labeled modules spanning immune, cell-cycle, oxidative-stress, and metabolic programs (Fig. 2C) and the top-ranked modules mapped to interferon signaling (module 42; n=114). Additional modules were enriched for proliferation (G2M Checkpoint / Mitotic Spindle; C5 module 1; n=85), oxidative-stress biology (Reactive Oxygen Species Pathway / Heme Metabolism; C5 module 12; n=82), and metabolic labeling (Oxidative Phosphorylation / Fatty Acid Metabolism; C5 module 4; n=77).

In the C2 network, the leading module was annotated by bile acid and xenobiotic metabolism (C2 module 4; n=56), alongside a prominent signaling/stress module (TGF Beta Signaling / UV Response DN; C2 module 1; n=43; Supplementary Table 10). Together, the C5 and C2 pathway networks show that coupling between ΔμG and ΔAging is not diffuse but instead concentrates into interpretable immune, proliferative, structural-remodeling, oxidative-stress, and metabolic modules. As a complementary gene-level concordance check, we performed rank-rank analysis (RRHO2)^22,23^ comparing ranked ΔμG and ΔAging gene-delta signatures (Supplementary Fig. 2).

### Gene-level network modules and metabolic-health genes anchor on concordant signals

To investigate the gene expression modules associated with specific pathway enrichments during longitudinal aging and simulated microgravity associated changes, we clustered genes by their across-donor coupling between ΔAging and ΔμG to define gene modules (Fig. 2E; Supplementary Table 11). The highest-ranking modules showed consistent positive coupling and mapped to coherent biological programs, including cell-cycle organization (mitotic spindle; module 1; n=448; ρ=0.454), inflammatory signaling (TNFα/NFκB; module 5; n=417; ρ=0.479), and innate immune effector biology (complement; module 3; n=372; ρ=0.526). Representative gene anchors within these modules included spindle-associated genes (NUMA1, PCNT, CEP57), canonical NFκB regulators (RELA, NFKBIA, TNFAIP3), and complement components (C3, C1QC), indicating that the subject-level changes are likely driven by immune and proliferative programs.

Beyond these core modules, the gene network also captured growth and nutrient-signaling structure aligned with the coupled response. For instance, the PI3K–AKT–mTOR module (module 4; n=246) exhibited strong positive correlation (ρ=0.543) and included pathway-relevant anchors such as PTEN and GRB2. Smaller modules captured inflammatory chemokine signaling (module 10; n=8; SERPINB2, CXCL1, IL1B) and MYC-linked programs (module 6; n=211), reinforcing that concordant ΔAging/ΔμG coupling organizes into a limited number of recognizable signaling and immune-growth programs (full module statistics in Supplementary Tables 3–6).

To provide concrete molecular anchors for metabolic concordance observed in the previous analyses (Fig 1B and C and Fig 2B), we focused on genes in the aging hallmarks metabolic-health category (Fig. 2F; Supplementary Table 12). Several showed strong positive coupling between ΔAging and ΔμG, including SLC25A1 (mitochondrial citrate/isocitrate carrier; ρ=0.697), SLC7A6 (amino-acid transport linked to nutrient sensing; ρ=0.696), and ALOX12 (lipid-peroxidation signaling; ρ=0.638).

These results suggest that substrate transport, mitochondrial carbon handling, and lipid/oxidation signaling are convergent nodes within the metabolic-health pillar.

### Vaccine-response signatures show concordant attenuation under aging and simulated μG

To test whether the shared immunometabolic remodeling extends to an immune-function–relevant program, we next examined vaccine-response gene signatures and their within-donor coupling under aging and simulated μG. To do so, we used previously defined gene expression profiles observed in humans after Day 1 and Day 3 post influenza vaccine administration ^24,25^. Both the day 1 and day 3 vaccine-response signatures showed negative adjusted contrasts for Aging vs Baseline and μG vs Baseline, consistent with reduced vaccine-response transcriptional activity under aging-related and simulated μG conditions after vaccination. The aging effect was evident on day 1 (estimate = −0.0293; p = 0.0334, Fig. 3A) and directionally consistent on day 3 (estimate = −0.0148; p = 0.212, Fig. 3D), with Fisher’s combined p = 0.042 across day 1/day 3 supporting an overall aging-associated attenuation of vaccine-response programs. μG effects were likewise negative in both datasets (day 1: estimate = −0.0133; p = 0.203, Fig. 3B; day 3: estimate = −0.0273; p = 0.0701, Fig. 3E), yielding a concordant trend on combination (Fisher p = 0.0749).

**Fig. 3.**
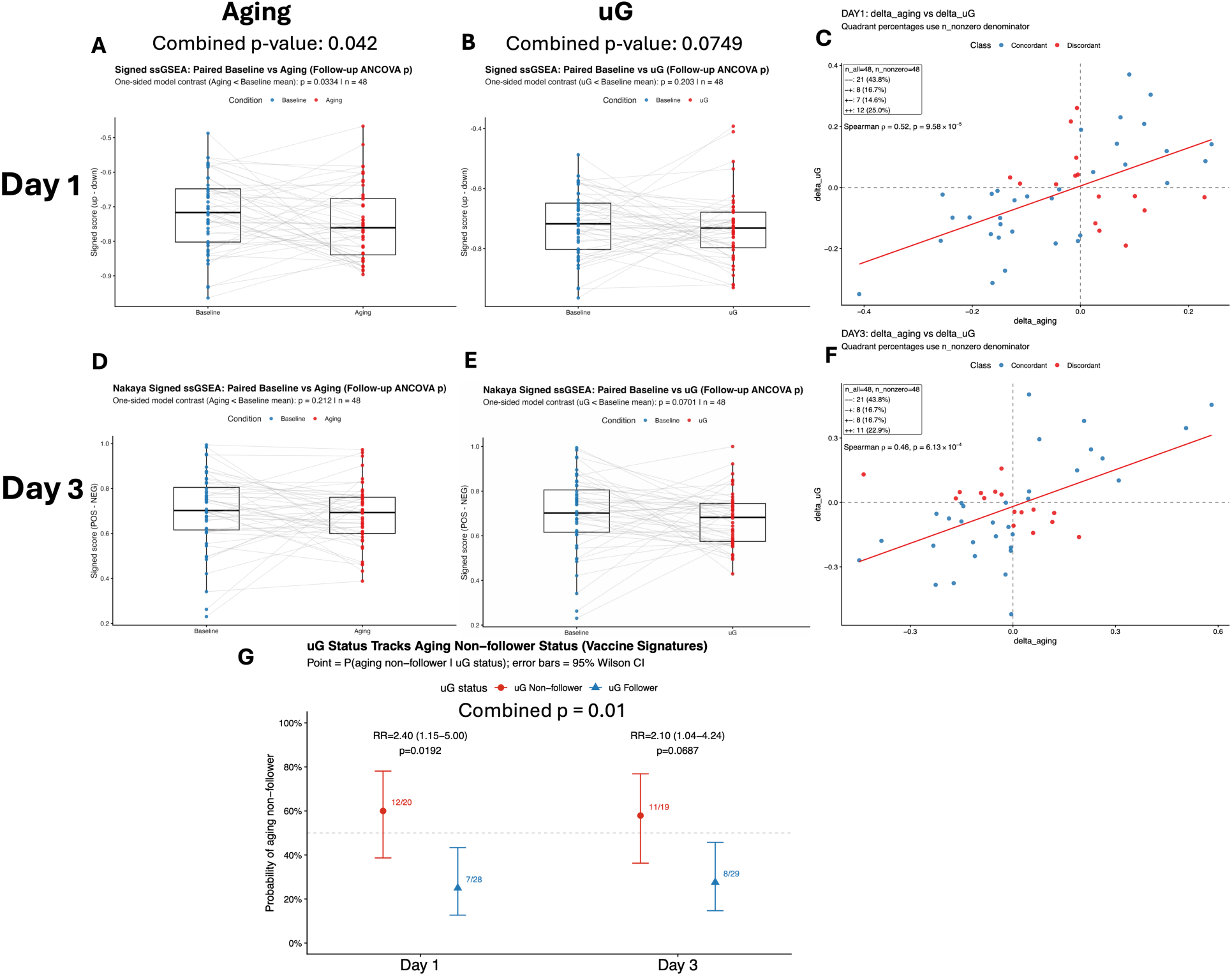
Vaccine-response signatures show concordant attenuation under aging and simulated microgravity. (A–B) Boxplots of per-donor signed vaccine-response ssGSEA scores for the day 1 signature, comparing Baseline vs Aging (A) and Baseline vs μG (B). (C) Donor-level concordance for the day 1 signature, plotting ΔAging (Aging − Baseline) versus ΔμG (μG − Baseline) across donors with fitted trend line; points are colored by sign-concordance class, and the panel annotates n_all, n_nonzero, quadrant percentages, and Spearman ρ with p-value. (D–E) Boxplots of per-donor signed vaccine-response ssGSEA scores for the published day 3 signature, comparing Baseline vs Aging (D) and Baseline vs μG (E). (F) Donor-level concordance for the day 3 signature, shown as in (C). (G) uG-to-aging tracking analysis: donors were classified as follower or non-follower based on whether delta-sign direction matched the cohort-direction sign from ANCOVA contrasts; points show *P*(aging non-follower ∣ uG status) with 95% Wilson confidence intervals, with day-specific risk ratios and Fisher’s exact p-values annotated.

At the individual level, donors showed consistent ordering across conditions with Δaging and ΔμG being positively correlated in both datasets (Spearman ρ = 0.519 and 0.457; p = 9.58×10⁻⁵ and 6.13×10⁻⁴, Fig. 3C,F). Despite the overall attenuation trend, donor responses were largely heterogeneous (Fig. 3C,F). Notably, simulated μG captured this heterogeneity in a way that aligned with longitudinal aging: donors who deviated from the cohort-average direction under μG were more than twice as likely to also deviate during aging (day 1: RR = 2.40, p = 0.0192; day 3: RR = 2.10, p = 0.0687; combined evidence across signatures p = 0.0101; Fig. 3G). These results indicate that μG recapitulates not only the cohort-average shift in vaccine-response transcriptional programs, but also a shared donor-stratifying response pattern.

These results show a large degree of variation in the vaccine response during aging and demonstrate that simulated microgravity exposure can at least in part, emulate this heterogeneity.

### SCENITH PBMC assays align aging and simulated μG energetic dependencies

To examine the immunometabolic effects of simulated microgravity and aging in an orthogonal functional context, we profiled energetic dependencies in PBMC using SCENITH, a flow cytometry-based method used to analyze the metabolic profiles at the single cell resolution, in young and older donors and under and simulated μG conditions (Fig. 4A).

**Fig. 4.**
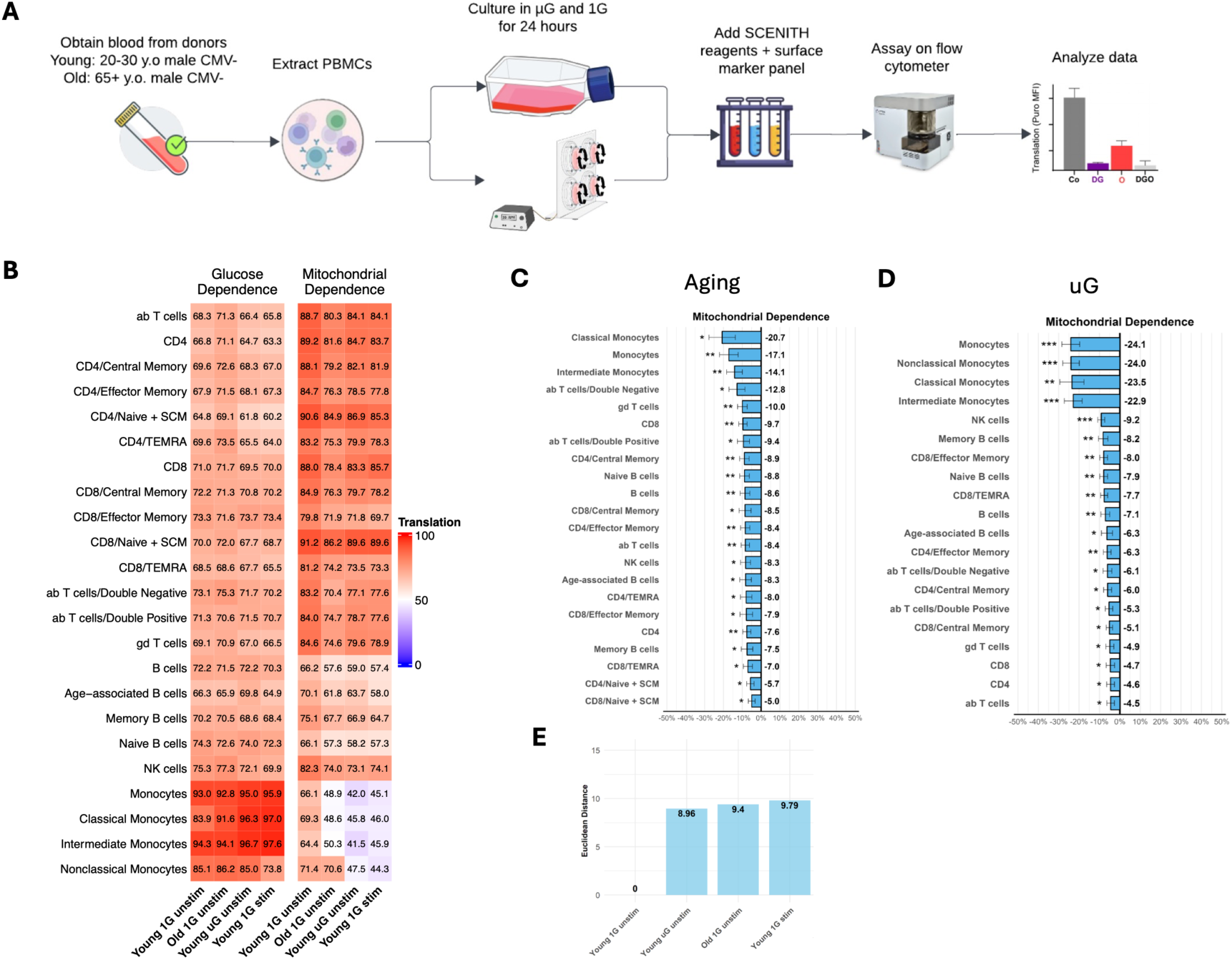
SCENITH functional energetics align aging and simulated microgravity dependency profiles across immune subsets. (A) SCENITH study design schematic. PBMCs from CMV-negative male donors (young: 20–30 years; old: 65+ years) were cultured under 1G or simulated μG for 24 hours; a subset received LPS stimulation during the final 6 hours. SCENITH was performed using puromycin incorporation with metabolic inhibitor conditions to derive glucose dependence and mitochondrial dependence across immune subsets. (B) Mitochondrial dependence across 23 PBMC subsets comparing Old 1G (aging-associated) versus Young 1G baseline. (C) Mitochondrial dependence across subsets comparing Young μG versus Young 1G baseline, and Old 1G versus Young μG, assessing similarity between aging-associated and μG-associated energetic states. (D) Glucose dependence across the same subsets/conditions, highlighting limited compensatory increases under aging or μG and subset-specific shifts under inflammatory stimulation. (E) Euclidean distance integration of glucose and mitochondrial dependence across subsets to summarize overall similarity among conditions and relate basal aging/μG profiles to an acute inflammatory (LPS) reference. Full dependency/capacity heatmaps and translation readouts are provided in Supplementary Figs. 4–6.

We found that aging was characterized by a broad shift away from mitochondrial dependence without a compensatory increase in glucose dependence (Fig. 4B–D). Specifically, 22/23 immune subsets show decreased mitochondrial dependence in Old 1G relative to Young 1G, while glucose dependence did not show a significant global increase. Monocytes were most significantly impacted by decreased mitochondrial dependence. Nonetheless, this pattern is consistent with a widespread reduction in mitochondrial reliance across immune subsets, consistent with the cohort-level concordance between longitudinal aging and simulated microgravity exposure. It sets the baseline for testing whether simulated μG elicits a similar energetic profile.

Simulated μG exposure yielded a closely aligned phenotype in young PBMCs (Fig. 4B–D). After 24 hours of μG, 20/23 subsets show decreased mitochondrial dependence, again without a significant increase in glucose dependence. Direct comparison of Old 1G and Young μG shows no statistical difference in 22/23 subsets, indicating strong similarity in energetic profiles. Together, these findings indicate a similar metabolic dependency profile in PBMCs under simulated μG.

Given that aging and μG showed overlapping metabolic profiles by the SCENITH assay, and that stimulation with TLR agonist can rewire immune cell metabolism, we next assessed how μG and aging SCENITH profiles change with TLR stimulation with LPS, which binds to TLR4. LPS stimulation yielded an acute inflammatory metabolic profile marked by reduced mitochondrial dependence across most subsets (18/23), with monocytes shifting toward aerobic glycolysis and NK cells showing reduced glucose dependence (Fig. 4B–D). Euclidean distance analyses further placed the basal aging and μG conditions near this inflammatory state: Young μG and Old 1G shift by approximately 9 units, comparable to the Young 1G LPS shift (9.79 units; Fig. 4E), while Old μG with stimulation shows a larger shift (15.36 units; Supplementary Fig. 3). These distances indicate that aging and simulated μG emulate energetic states with LPS-like profile. This pattern aligns with the broader mitochondrial/oxidative-stress coupling observed in pathway-level analyses and provides a functional anchor for those transcriptomic themes. Full dependency and capacity heatmaps and translation readouts are provided in Supplementary Fig. 4–6.

Together, these findings provide a functional readout validating the findings observed in gene expression both at the level of DiseaseAge for the metabolic system and the aging hallmark for metabolic health profiles (Fig. 2).

## Discussion

Here, we find that individual responses to simulated microgravity closely track longitudinal aging trajectories across a multi-scale biological landscape. This concordance extends from system-level pillars to gene networks, vaccine response profiles and functional energetics. The pattern is consistent with broad, donor-coupled remodeling rather than isolated pathway perturbations. By employing aging as a comparative lens, we resolve this complex response into coordinated themes of mitochondrial stress, immunometabolism, and cytoskeletal reorganization ^26,27^.

At the system scale, the robust coupling across aging hallmarks and DiseaseAge domains establishes a cohort-wide alignment that bridges phenotypic risk to molecular mechanisms. This finding is most notable in the metabolic and musculoskeletal categories but extends to multiple biological aging features. A global positive shift in pathway-level concordance reinforces this systemic signal and organizes it into coherent network modules rather than a single driver. While broad metabolic themes in Gene Ontology and canonical pathways show strong alignment, the specificity of the signal points to a stress-response mechanism rather than a simple upregulation of metabolic machinery. Specifically, the data highlight mitochondrial quality control and ROS handling over general Hallmark respiration ^28,29^. This interpretation is supported by our gene-level analysis where anchors such as SLC25A1 and ALOX12 link mitochondrial transport and lipid peroxidation to the donor-level aging shift. Consistent with this pattern, the concordance also extends beyond pathway structure to an immune-function–relevant program: a targeted vaccine-response transcriptional program also showed concordant attenuation under both aging and simulated μG. Vaccine-response signatures were reduced at follow-up, and donors with larger aging-associated decreases tended to exhibit larger μG-associated decreases, indicating shared donor-specific remodeling in an immune-function–relevant axis.

An important feature of these vaccine-response programs was inter-individual variability. A substantial subset of participants (1/3) changed in a different direction to the cohort-average trend. We and others have shown that this is a common behavior when analyzing longitudinal vs cross-sectional human data ^30^. Notably, simulated μG recapitulated this structure, as individuals who deviated under μG were more likely to deviate during aging as well (risk ratios >2) (Fig. 3G). Together, these findings suggest that simulated microgravity can serve as an experimentally tractable perturbation to reveal donor-stratifying immune-response remodeling linked to longitudinal aging, complementing cohort-average analyses and highlighting differences in susceptibility or resilience relevant to vaccination.

This robust alignment was also reflected in functional readouts, spanning an immune-response program (vaccine-response signatures) and cellular energetics (SCENITH). Our SCENITH profiling in an independent cohort provides orthogonal validation by revealing that both aging and simulated microgravity induce a specific energetic phenotype characterized by a widespread reduction in mitochondrial dependence without a compensatory glycolytic switch ^17,29^. This severely restricted metabolic flexibility mirrors recent SCENITH-based energetic profiling of aged human immune cells, which identified a similar coupling of impaired metabolic flexibility with hyperinflammatory responses ^31^. Such energetic inflexibility directly contrasts with the classical Warburg effect often seen in activation and places the microgravity-associated phenotype closer to a state of energetic inflexibility or exhaustion. Quantitatively, this state aligns with an acute LPS-induced inflammatory profile and reinforces the biological plausibility of the immunometabolic and oxidative-stress themes identified in our pathway analysis. Such energetic inflexibility directly impedes the rapid metabolic reprogramming, specifically the upregulation of oxidative phosphorylation and glycolysis, required to fuel successful vaccine responses ^26,27^. Consequently, the profound energetic constraints induced by simulated μG mimic the mitochondrial dysfunction known to blunt vaccine efficacy in older adults ^32–34^. Furthermore, the robust alignment of ROS pathways alongside interferon, TNF/NFKB, and complement modules suggests that this energetic constraint may be intrinsically linked to innate immune priming or sterile inflammation, creating a noisy, hyper-reactive baseline state that drowns out productive vaccine-induced transcriptional programs ^26,34,35^.

Our results also highlight a potential structural basis for these shifts. The strong concordance in mechanotransduction and cytoskeletal pathways suggests that physical unloading may act as the upstream trigger that shapes immune-cell state and energetic demand ^36,37^. While the direct causal link remains to be tested, the coherent modular organization of these pathways implies that cytoskeletal remodeling is not an isolated adaptation but is intimately coupled to the broader stress response, including potentially mitochondrial dysfunction.

A primary strength of this study is the paired, within-donor design. This approach allows us to quantify the alignment of microgravity and aging shifts while controlling for the high inter-individual variability inherent in human cohorts. The integration of orthogonal energetic assays further strengthens the biological interpretability of the transcriptomic correlations. However, several limitations warrant consideration. First, the SCENITH assays were restricted to PBMCs from CMV-negative male donors, which limits generalization to other tissues, sexes, or viral histories. Second, the 24-hour duration of simulated microgravity captures acute adaptation which may differ from chronic in-flight responses. Finally, while the coupling is robust and statistically significant across scales, it remains correlational. Future work is required to establish the directionality of these relationships and whether they represent maladaptive stress or successful adaptation.

The multi-scale convergence of microgravity and aging signatures suggests that simulated microgravity is a practical and high-fidelity platform for modeling aspects of human biological aging ^10,38^. The specific identification of mitochondrial stress and energetic constraint as central features provides a targeted rationale for countermeasure development. Rather than broad anti-inflammatory or metabolic interventions, strategies aimed specifically at preserving mitochondrial respiratory capacity and energetic resilience may be effective at decoupling spaceflight stress from aging-like degeneration. Lastly, these findings establish simulated microgravity as a tractable platform for modeling accelerated aging, supporting a mechanism where mitochondrial, ROS, and immunometabolic programs are co-modulated under shared stress conditions ^26,27^.

## Methods

### Study design and cohorts

We employed a within-donor paired design to analyze 48 donors who had high-quality RNA-sequencing profiles available across three distinct conditions: Baseline (1G; standard gravity), Baseline (μG; simulated microgravity), and Follow-up (natural aging at 1G). Participants were aged 24–90 years at baseline. Each participant contributed paired samples across all three states, enabling within-individual contrasts that minimize inter-person variability. The median interval between Baseline and Follow-up was 6.96 years (IQR 5.51–8.37; min 4.03; max 9.27).

Deidentified cryopreserved peripheral blood mononuclear cell (PBMC) samples were profiled by bulk RNA-seq, with primary contrasts defined as ΔμG = Baseline(μG) - Baseline(1G) and ΔAging = Aging(1G follow-up) - Baseline(1G), computed within each donor. A separate PBMC cohort was used for SCENITH-based metabolic profiling under 1G vs simulated μG culture. Full cohort details and protocol are provided in *SCENITH-based metabolic profiling*.

#### Ethics statement

All human samples were obtained as deidentified specimens from 1KIP cohort, in accordance with Buck Institute policies and under IRB approval or an IRB-exempt determination as applicable. Written informed consent was obtained by the originating study, and only deidentified data were analyzed.

### Human blood sample and cell culture

Cryopreserved PBMCs from 1KIP cohort were thawed, washed, and counted for viability, then resuspended at 1 × 10^6 cells/mL in complete RPMI-1640 medium (10% FBS, 2 mM L-glutamine, 1% penicillin/streptomycin, 0.1 mM non-essential amino acids, 1 mM sodium pyruvate, 10 mM HEPES, 50 μM 2-mercaptoethanol) and allowed to recover for 1 hour at 37°C and 5% CO2. For each donor, cells were then split into 1G and simulated microgravity (μG) conditions and cultured for 24 hours at 37°C and 5% CO2. The 1G condition was maintained in standard static tissue culture plates or flasks, and the μG condition was maintained in 2 mL rotating wall vessels (RWV) within an RCCS-4D bioreactor at 15 RPM ^4,39^.

### RNA-seq preprocessing and normalization

Total RNA was extracted using RNeasy Plus Mini Kit (Cat# 74134, Qiagen) as per the manufacturer’s instructions. RNA quantity check, preparation of RNA library, and mRNA sequencing were conducted by Novogene Co., LTD (CA, US). About 20 million paired-end 150 bp reads per sample were generated on the Illumina NovaSeq X Plus Sequencing System. FASTQ raw reads were analyzed using the nf-core/rnaseq pipeline ^40^ and produced a gene expression matrix.

Raw gene-by-sample count matrix were harmonized with sample metadata (Supplementary Table 14). Counts were normalized with the trimmed mean of M-values (TMM) method implemented in edgeR ^41,42^ and transformed to logCPM with a prior count. Genes were filtered to retain CPM > 1 in at least 10 % of samples. Batch correction used ComBat from the sva package ^43^, and covariates Sex and Age were included when the design matrix was full rank. When an external reference cohort was included, ComBat was anchored to the reference batch (GTEx) ^19^.

### Aging hallmark predictive models

We developed predictive models for 14 hallmarks of aging, which we divide into protectors and drivers. We define aging protectors as the biological processes that enhance our defenses against aging (intrinsic capacity and resilience), and aging drivers as the biological processes that fuel aging (transcriptomic age, chronic inflammation, epigenetics, immune health, intracellular communication, metabolic health, mitochondria, nutrient sensing, proteostasis, senescence, stem cell exhaustion, telomere).

Predictive models for most pillars were built using elastic-net regularized generalized linear models implemented in the R package glmnet ^44,45^. We used whole-blood RNA-seq data from the GTEx ^19^ population reference and the pillar gene sets to build age predictors. For intrinsic capacity, we used genes previously identified as associated with the DNAm Intrinsic Capacity clock ^20^. For resilience, we used a set of genes associated with DNA damage, stress response, and autophagy ^46,47^. For senescence and stem cell exhaustion, we used a curated gene set identified in multiple publications ^48–54^. For chronic inflammation, we used MSigDB and extracted inflammation-related pathway genes. For intracellular communication, we extracted from MSigDB genes involved in signal transmission across the cell membrane via G-protein-coupled receptor activity. For nutrient sensing, we used MSigDB genes involved in the regulation of triglyceride sequestration. For proteostasis - MSigDB genes involved in regulating the unfolded protein response in the endoplasmic reticulum. For epigenetics - MSigDB genes involved in epigenetic regulation of gene expression. For telomere - MSigDB genes involved in protein localization to telomeric regions of chromosomes. For mitochondria - MSigDB genes involved in the aggregation and arrangement of the mitochondrial respiratory chain complex. A predictive model for metabolic health was built in 2 steps, incorporating metabolic flux calculations via the METAFlux R package ^55^. In the first step, we fit an elastic-net model to predict age from the metabolic reaction corpus. In the next step, we built an elastic-net model to predict the resulting Age residuals from enzyme expression in the reactions selected in the first step.

A predictive model for immune health was also built in 2 steps. In the first step, we used the OneK1K ^56^ reference dataset and fit an elastic-net model to predict cell proportions from the gene expressions in the corresponding cell types. In the second step, we built an elastic-net model to predict individuals’ ages from cell proportions.

### Aging hallmark and DiseaseAge scoring and correlation analysis

We quantified biological aging using two complementary scoring approaches. BeyondAge pillar scores were computed locally from ComBat-adjusted logCPM values using a pre-defined coefficient matrix derived from aging hallmark-based models and anchored to a GTEx population reference (n = 803). For each pillar, a weighted sum of member genes was calculated. Pillars with fewer than three mapped genes were assigned missing values. In parallel, DiseaseAge predictions were obtained from an external API/ website ^18^ and processed using the same donor-aggregation strategy.

For both methods, scores were aggregated to per-donor, per-condition means. We then computed within-donor deltas for each pillar and disease domain: ΔAging (Aging 1G - Baseline 1G) and ΔμG (Baseline μG - Baseline 1G). Pearson correlations between ΔAging and ΔμG were calculated using complete cases. To test for positive coupling, we utilized one-sided p-values (alternative = greater) for nominal significance labeling, while two-sided 95% confidence intervals were used to quantify effect-size uncertainty. A combined significance value across pillars or domains was computed using Fisher’s sumlog method ^57^.

### Prediction modeling

Predictive analyses were performed separately for each BeyondAge pillar and DiseaseAge domain/system using univariate linear regression (ΔAging = β0 + β1·ΔμG) on complete-case donors. Performance was estimated by leave-one-out cross-validation (LOOCV), in which each donor was held out once (number of folds = number of donors for that feature), the model was fit on the remaining donors, and an out-of-fold prediction was generated for the held-out donor. MAE, RMSE, and cross-validated R² were calculated from these out-of-fold predictions. Accuracy was summarized as 1 − (MAE / ΔAge Range), where ΔAge Range = max(ΔAging) − min(ΔAging) within each pillar/domain.

### ssGSEA pathway scoring and ΔAging/ΔμG correlation analysis

ssGSEA^21^ was performed on ComBat-adjusted logCPM expression values with sample metadata to define three conditions (Baseline, Aging, μG). Gene symbols were used as identifiers and duplicated symbols were collapsed by mean expression. Donor matching across conditions was enforced to enable paired within-donor comparisons. Gene sets were obtained from MSigDB via msigdbr ^58^, including Hallmark (H), C2, and C5 (BP/CC/MF) collections ^59,60^, and were restricted to genes present in the expression matrix. ssGSEA scoring was carried out using the GSVA implementation of ssGSEA with normalized scores ^16^, retaining a consistent pathway universe across conditions.

For each pathway, Spearman correlations between ΔAging and ΔμG were computed across donors (minimum n = 3 donors with complete data) and adjusted for multiple testing using the Benjamini–Hochberg procedure ^61^; the number of contributing donors was recorded. Global concordance was summarized by density distributions of pathway-wise correlations with a zero-reference line; because pathways are not statistically independent, a one-sided Wilcoxon signed-rank test against zero was used only as a descriptive summary of distribution direction.

### Theme-level pathway summaries

Pathway-level correlation tables from Hallmark, C2:CP, and C5:GO collections were merged to generate theme summaries. Biological themes (mechanotransduction, cytoskeleton, mitochondria, ROS, metabolism) were summarized using curated keyword filters that map pathways to each category. The theme-matched summary table (Supplementary Table 7) includes pathways passing nominal significance (p < 0.05). For figure highlights, top five pathways per theme were selected as the highest Spearman correlation coefficient among significant theme-matched pathways.

### Pathway similarity networks

Pathway similarity networks were constructed separately for MSigDB Hallmark (H), C2, and C5 collections using the pathway-wise ΔAging/ΔμG correlation results from the ssGSEA analysis. Nodes were defined as pathways present in the corresponding MSigDB collection that passed a nominal association filter (p < 0.05). Each node was annotated with its correlation statistic (ρ), nominal p-value, FDR-adjusted p-value, and gene-set size. Edges represented gene-set overlap quantified by Jaccard similarity, computed using a sparse gene-by-pathway incidence representation; only non-zero overlaps were retained.

To obtain comparable network density across collections, a collection-specific Jaccard threshold was selected to target an expected average node degree of 12. Thresholds were determined by sampling up to 50,000 random pathway pairs, estimating the non-zero Jaccard distribution, and selecting a high quantile as the cutoff; thresholds and diagnostics were recorded. Undirected weighted graphs were then constructed using Jaccard values as edge weights, and community structure was identified by weighted Louvain clustering ^62^. For visualization, edges were pruned for readability (default: top 25% of edge weights when sufficient edges were available), isolates were removed, and the largest connected component was plotted using a Fruchterman–Reingold layout ^63^. Node size was scaled by |ρ| and node color indicated module membership.

Module labels were assigned in two complementary ways. First, within each module, representative pathways were identified by weighted node strength after removing collection-specific prefixes (e.g., GOBP_, REACTOME_, HALLMARK_). Second, to provide a compact and interpretable vocabulary for C2 and C5 modules, modules were labeled by Hallmark enrichment: Hallmark gene sets were tested for enrichment among the pathways in each module using a hypergeometric test with Benjamini–Hochberg correction. Module–Hallmark scores were computed as the sum of pathway contributions (strength × −log10(FDR)) and normalized by total module strength, and the top two Hallmarks were used as module labels.

### Gene-level correlations and gene network analysis

Gene-level delta matrices were computed from the logCPM-normalized, batch-corrected expression matrix described in “RNA-seq preprocessing and normalization.” Samples were aligned to metadata and matched across Baseline, Aging, and μG conditions with consistent donor ordering. For each gene, within-donor deltas were computed (ΔAging and ΔμG), and coupling between ΔAging and ΔμG was quantified across donors using Spearman correlation (minimum n = 3 donors with complete data), with Benjamini–Hochberg correction and reporting of the number of contributing donors per gene.

For gene network construction, genes passing a nominal association filter (p < 0.05) were retained and used to compute gene–gene similarity based on their combined delta profiles. For each gene, ΔAging and ΔμG vectors were concatenated across donors, rank-transformed (average ranks for ties), and z-scored across donors; genes with zero variance or non-finite values were removed. Pairwise similarity between genes was calculated as the normalized dot product of these scaled rank profiles. To obtain comparable network density, a similarity threshold was selected by sampling up to 50,000 random gene pairs to target an expected average node degree of 12 (threshold constrained to 0.1–0.9), after which edges exceeding the threshold were computed in blocks to enable efficient construction. The resulting undirected weighted graph was summarized (nodes, edges, density, largest connected component) and partitioned into modules using weighted Louvain community detection.

Modules were filtered to retain those with at least four genes. For visualization, isolates were removed and the largest connected component was plotted using a Fruchterman–Reingold layout with edges pruned for readability (top 10% by weight); node size reflected |ρ| for the gene-level ΔAging/ΔμG coupling and node color indicated module membership. Module-level behavior was summarized by computing module scores as the mean delta across genes within each module for Aging and μG, and by correlating module scores across donors. Functional interpretation used Hallmark enrichment of module gene sets using a hypergeometric test with Benjamini–Hochberg correction (MSigDB Hallmark sets via msigdbr, with a GMT fallback when needed), and top enriched Hallmark terms per module were reported. For metabolic-health pillar gene anchors, genes within the BeyondAge metabolic-health pillar were analyzed individually by correlating ΔAging and ΔμG across donors and reporting effect sizes with two-sided confidence intervals.

### Vaccine-response signature scoring and follow-up modeling

Donor-level signed ssGSEA scores were computed for two independent published vaccine-response gene signatures: a day 1 signature ^24^ and a day 3 signature ^25^. For each donor and condition (Baseline, Aging, μG), ssGSEA was performed separately for directionally opposing gene sets. A signed score was defined as the enrichment score for the pro-response set minus the enrichment score for the anti-response set, such that higher values indicate a stronger pro–vaccine-response transcriptional state.

To test follow-up effects while accounting for donor-specific starting levels, we fit a baseline-adjusted mixed ANCOVA model separately for each signature using a donor random intercept. The outcome was the signed follow-up score, and predictors included follow-up condition (Aging or μG), the donor’s baseline signed score (mean-centered across donors), and their interaction. Contrasts were tested for Aging and μG relative to the baseline mean level implied by the centered baseline covariate (Supplementary Table 13).

To assess whether donors show similar response magnitudes under aging and simulated μG, we computed donor-specific deltas (ΔAging and ΔμG) and tested Spearman concordance between ΔAging and ΔμG for each signature. p-values for model contrasts were one-sided (testing for reduced vaccine-response activity at follow-up). Evidence across the day 1 and day 3 signatures was summarized using Fisher’s combined p-values.

### Vaccine-response heterogeneity and cross-condition tracking analysis

To assess inter-individual heterogeneity beyond cohort-average shifts, we evaluated whether each donor’s within-donor change in vaccine-response signature followed the dominant direction observed at the cohort level. For each vaccine-response signature (day 1 and day 3), we used the sign of the cohort-level effect (estimated from the paired follow-up model) as the reference direction. Donors were classified as “followers” if their within-donor delta (follow-up minus baseline) matched the cohort-level direction, and as “non-followers” if their delta was opposite.

To test whether heterogeneity was shared across conditions, we quantified whether μG non-follower status was associated with aging non-follower status using 2×2 contingency tables. We summarized association using the risk ratio (RR) comparing the probability of being an aging non-follower among μG non-followers versus μG followers, and evaluated significance using Fisher’s exact test. Evidence across day 1 and day 3 signatures was additionally summarized using Fisher’s method. Confidence intervals for conditional probabilities displayed in Fig. 3G were computed using Wilson intervals.

### SCENITH-based metabolic profiling and analysis

PBMCs were obtained from concentrated buffy coats of CMV-negative male donors (young: 20–30 years, n=10; old: ≥65 years, n=10) and processed within 24 hours of collection. PBMCs were isolated by Ficoll-Paque density separation, assessed for viability, and cultured at 1×10^6 cells/mL in complete RPMI-1640 (2.05 mM L-glutamine, 10% FBS, 1% penicillin/streptomycin, 1% non-essential amino acids, 1% sodium pyruvate, 1% HEPES, 0.1% 2-mercaptoethanol). For each donor, cells were split and cultured for 24 hours at 37°C/5% CO₂ under 1G (tissue-culture flasks) or simulated microgravity in 10 mL rotating wall vessels (RCCS-4D bioreactor, 15 rpm). After incubation, cells were harvested, re-counted for viability, and plated at 5×10^6–1×10^7 cells/mL in 96-well U- or V-bottom plates for SCENITH.

SCENITH was performed using the classical kit (GammaOmics) following Argüello et al.^17^, with puromycin incorporation as a proxy for protein synthesis/ATP demand. Reagents were pre-warmed (37°C, 30 min), and cells were treated in parallel with DMSO control; 2-deoxy-D-glucose (100 mM, 15 min); oligomycin A (1 µM, 15 min); sequential 2-deoxy-D-glucose then oligomycin (10 + 5 min); harringtonine (2 µg/mL, 15 min); and puromycin (10 µg/mL) was added during the final 30 min. Cells were washed with ice-cold FACS buffer, Fc-blocked, stained with LIVE/DEAD dye and surface antibodies, fixed/permeabilized (Foxp3/Transcription Factor buffer set), and stained intracellularly with anti-puromycin (Alexa Fluor 647) before acquisition on an Aurora spectral flow cytometer. PBMCs were gated into 23 immune cell subsets using the predefined gating strategy (Supplementary Fig. 7), and SCENITH dependencies/capacities were computed using the standard SCENITH equations (Supplementary Fig. 8). For inflammatory stimulation, LPS was added after 18 hours of culture to 100 ng/mL and maintained for 6 hours (24 hours total) prior to SCENITH processing as above.

For statistical testing, pairwise group comparisons were performed per immune subset and metric using two-tailed Welch’s t-tests, with Benjamini–Hochberg FDR correction across all measured cell subsets (FDR < 0.05 considered significant). To summarize whole-profile similarity between conditions, Euclidean distance was computed in two-dimensional metabolic space defined by normalized glucose dependence (x-axis) and normalized mitochondrial dependence (y-axis): for each condition, a centroid was calculated as the mean of these normalized dependence values across the 23 subsets, and distances were computed relative to the Young 1G unstimulated centroid as d = sqrt((x_group − x_ref)^2 + (y_group − y_ref)^2).

## Data availability

Raw RNA-seq data generated in this study will be deposited in the Gene Expression Omnibus (GEO) prior to journal publication.

## Code availability

All code used for data processing, statistical analysis, and figure generation is available in a public GitHub repository: https://github.com/FEI38750/μG_PBMC

## Acknowledgements

We would like to thank BioRender for providing the tools used to create the figures included in this manuscript.

## Author contributions

F.W., A.C., D.A.W., and D.F. conceived the study and designed the experimental and analytical strategy. F.W. and A.C. led and performed the experimental work, with support from K.S. and A.S. F.W. led the computational and statistical analyses. A.R. provided the aging hallmark algorithms and analytical framework. M.F. contributed the DiseaseAge algorithm. K.S. supported additional analyses and interpretation. H.H, N.B. and M.D. provided samples and cohort access. F.W. and A.C. wrote the original draft. All authors reviewed and edited the manuscript and approved the final version. D.F. and D.A.W. supervised the work and secured funding.

