## Supplemental Figures for "Simulated Microgravity Recapitulates Aspects of Biological Aging in Humans"

### Supplementary Fig. 1 (related to Fig.1)

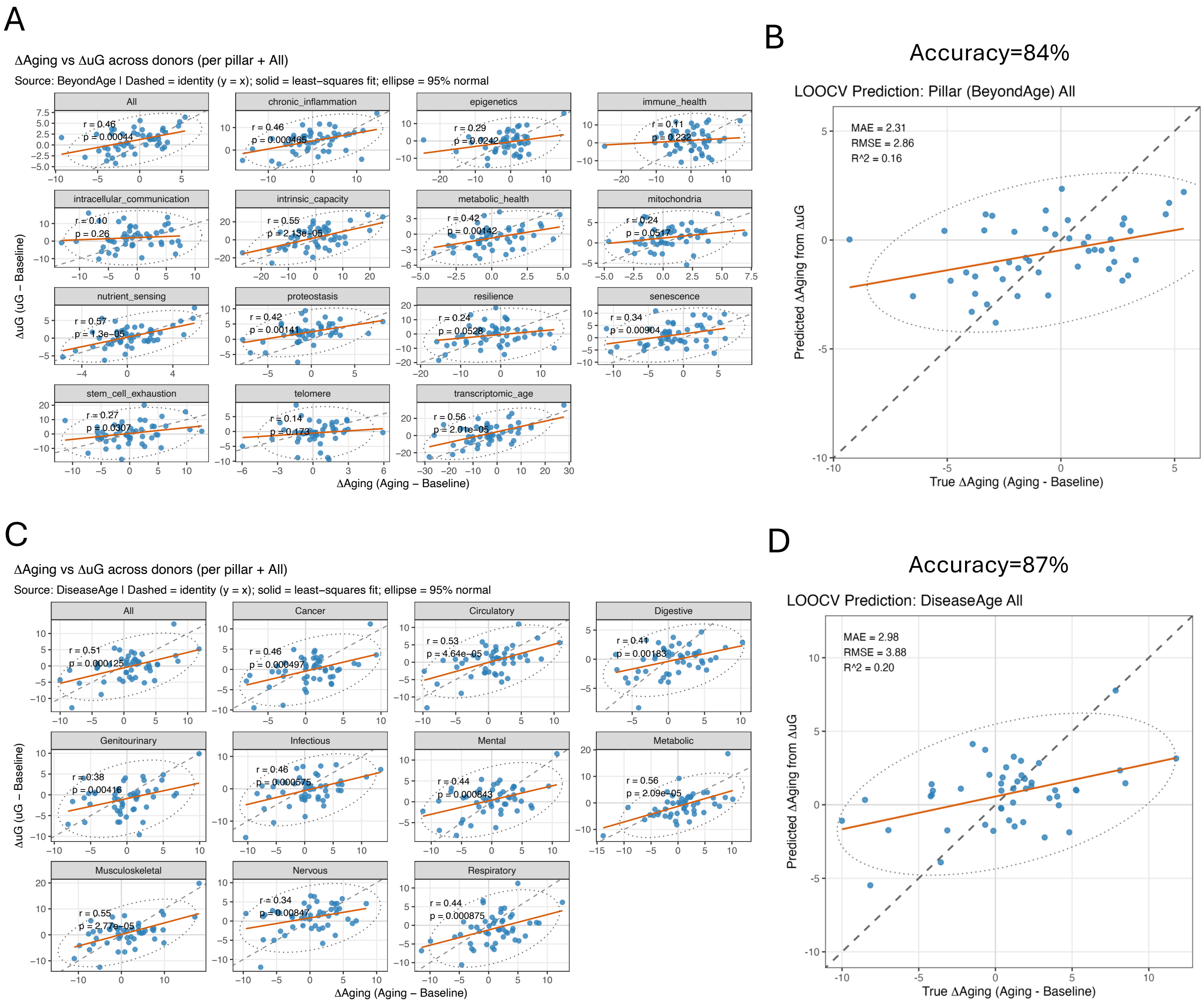


**Supplementary Fig. 1.**Donor-level Delta Aging vs Delta uG relationships and All-aggregate prediction scatter for BeyondAge and DiseaseAge. (A) Faceted donor-level scatter plots for BeyondAge pillars, showing Delta Aging vs Delta uG with an identity line, least-squares fit, 95% normal ellipse, and panel-wise Pearson annotations. (B) BeyondAge All-aggregate prediction scatter of observed vs LOOCV-predicted Delta Aging, with identity line and fitted trend; MAE, RMSE, and cross-validated R2 are annotated in-panel. (C): Faceted donor-level scatter plots for DiseaseAge domains using the same conventions as in (A). (D): DiseaseAge All-aggregate prediction scatter using the same conventions as in (B). These panels provide donor-level context for the correlation summaries in Fig. 1B-C.

Supplementary Fig. 2 (related to Fig.1-2)

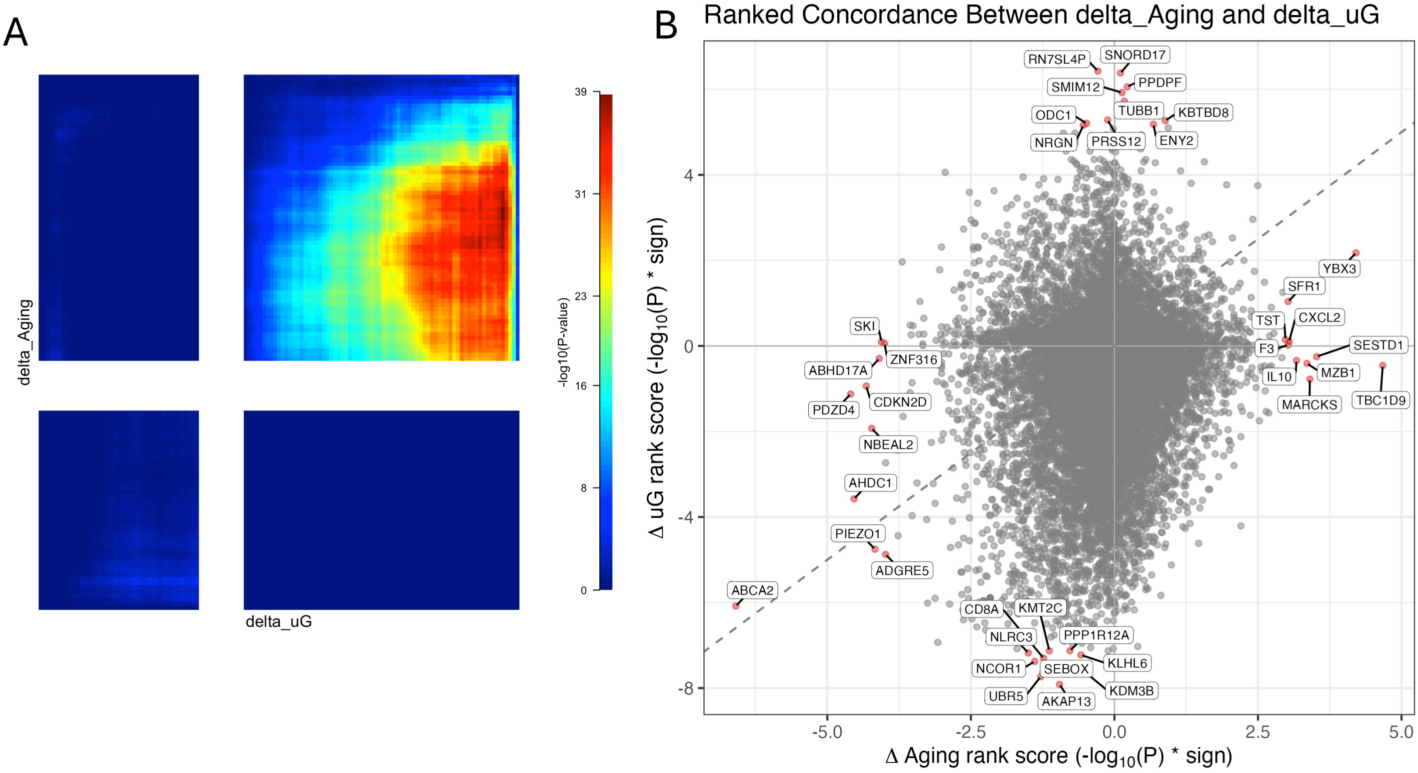


**Supplementary Fig. 2.** RRHO2 analysis of gene-delta signatures between microgravity and aging. (A) RRHO2 heatmap comparing ranked gene-level delta signals for ΔuG (uG − Baseline) and ΔAging (Aging − Baseline). Genes were ranked by signed Wilcoxon score (−log10(P) × sign(median Δ)), and heatmap intensity shows overlap significance (−log10 BH-adjusted P).
(B) Scatter plot of the same ranked scores for shared gene symbols (x-axis: ΔAging, y-axis: ΔuG). The dashed diagonal marks concordance between conditions; opposite quadrants indicate discordant directionality. Labeled points denote top extreme-ranked genes.

Supplementary Fig. 3 (related to Fig.4)

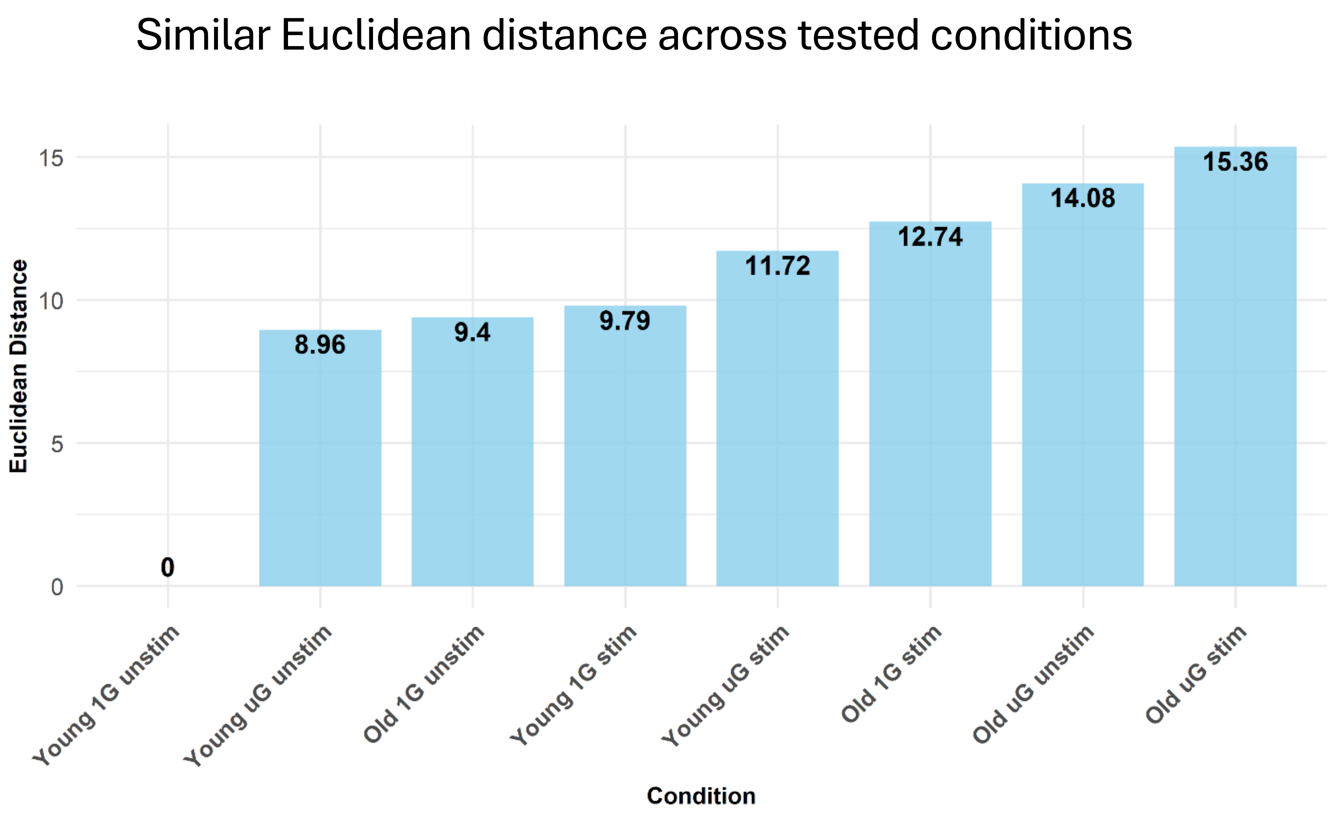


**Supplementary Fig. 3**: Euclidean distance analysis of overall similarity between the metabolic profiles of all stimulated and unstimulated experimental groups. The minimal distance between unstimulated Old 1G and Young μG groups and stimulated Young 1G group, contrasted with the large distances of these groups from the Young 1G baseline, and an even larger distance of remaining groups from the Young 1G baseline. An adjusted p-value (FDR) of less than 0.05 was considered statistically significant.

Supplementary Fig. 4 (related to Fig.4)

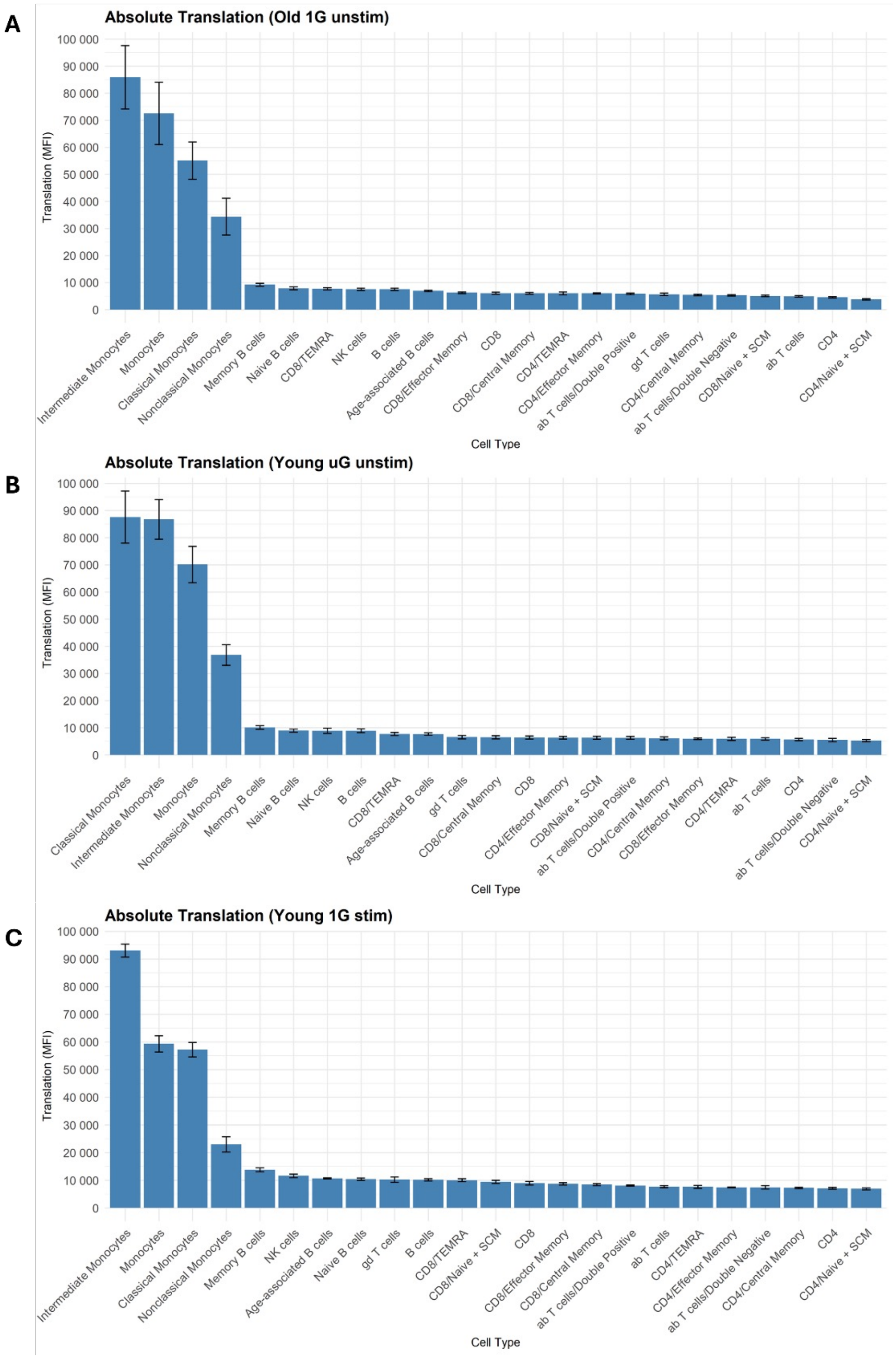


**Supplementary Fig. 4**: Absolute translation in aged, microgravity-exposed, and stimulated cells. Absolute translation measured in the geometric mean fluorescence intensity of the anti-puromycin antibody in (A) Old 1G unstim (aged), (B) Young μG unstim (μG-exposed), and (C) Young 1G stim (stimulated with LPS) PBMCs.

Supplementary Fig. 5 (related to Fig.4)

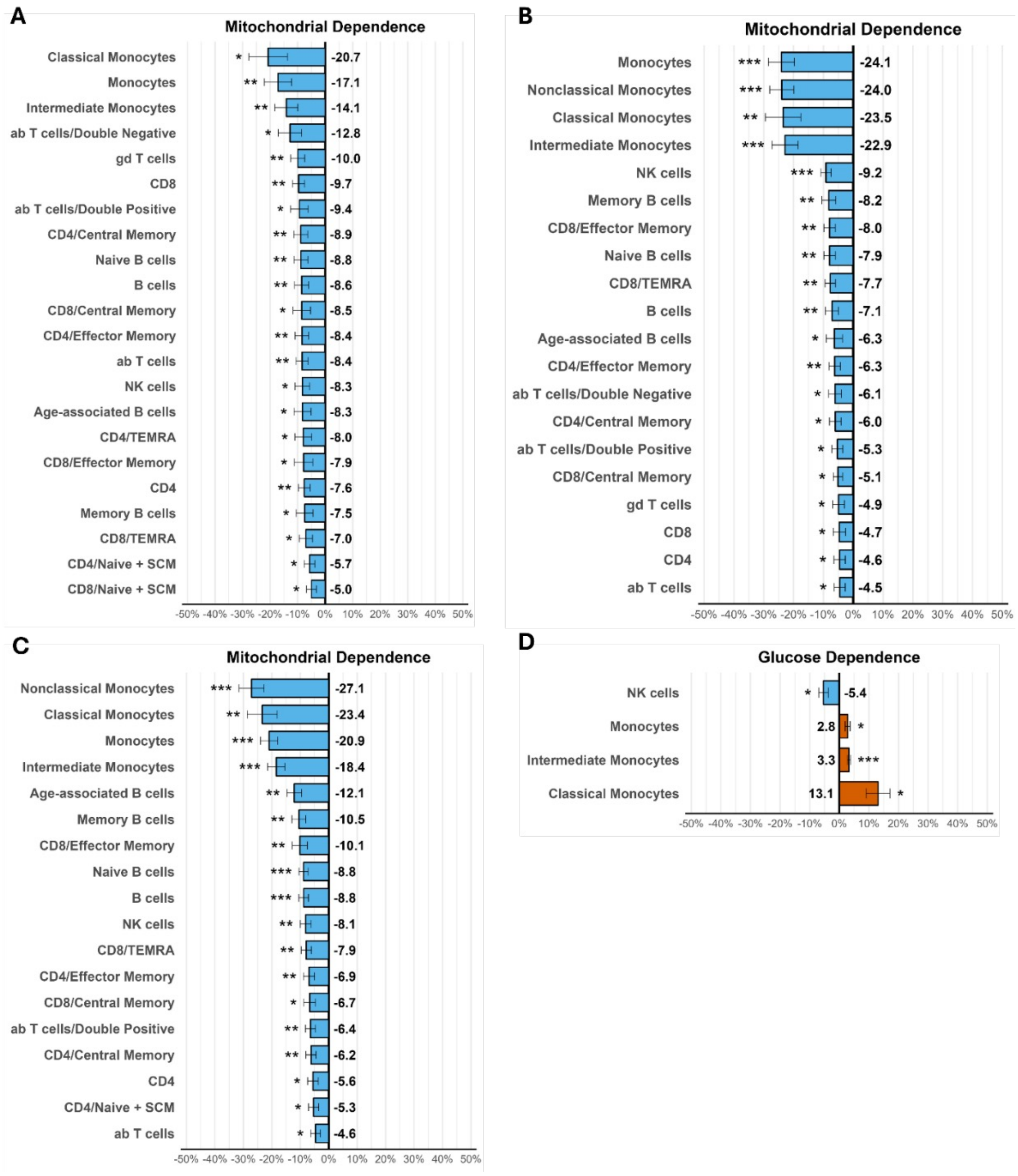


**Supplementary Fig. 5**: Statistically significant differences between metabolic profiles of different stressors compared to the baseline. Bar charts of statistically significant differences in mitochondrial dependence and glucose dependence of different immune cell subsets compared to the Young 1G unstim (baseline). Highest reduction in mitochondrial dependence was observed in monocytes in all three conditions. (A) Statistically significant reduction in mitochondrial dependence in 22 out of 23 PBMCs subtypes in the Old 1G unstim group. (B) Statistically significant reduction in mitochondrial dependence in 20 out of 23 PBMC subtypes in the Young μG unstim group (C) Statistically significant reduction in mitochondrial dependence in 18 out of 23 PBMC subtypes in the Young 1G stim group. (D) Statistically significant increase in glucose dependence in monocytes and reduction in NK cells in the Young 1G stim group.

Supplementary Fig. 6 (related to Fig.4)

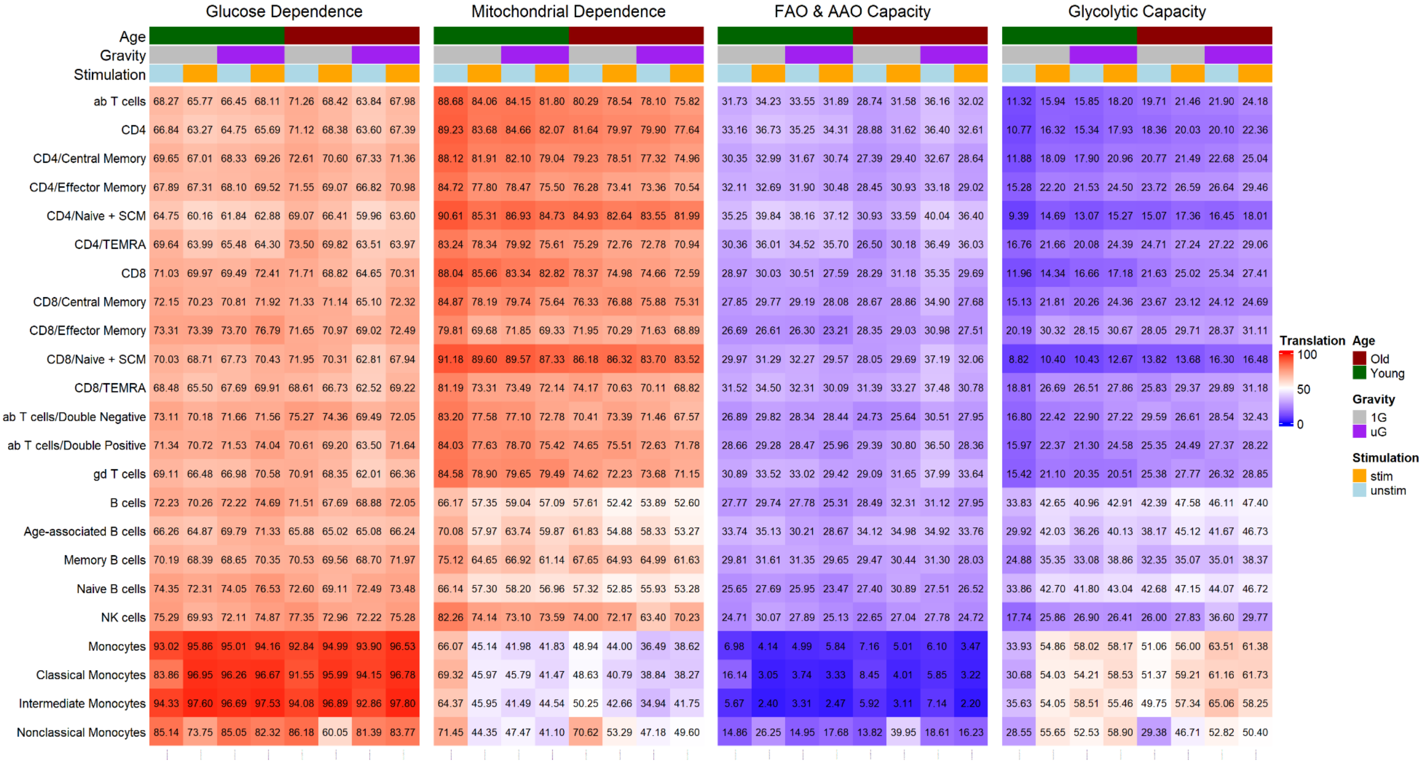


**Supplementary Fig. 6**: Heatmap of all SCENITH capacities and dependencies. Full heatmap including all conditions (age, gravity, stimulation) and metabolic profiles (glucose dependence, mitochondrial dependence, FAO & AAO capacity, glycolytic capacity). Capacities are derived parameters.

Supplementary Fig. 7 (related to Fig.4)

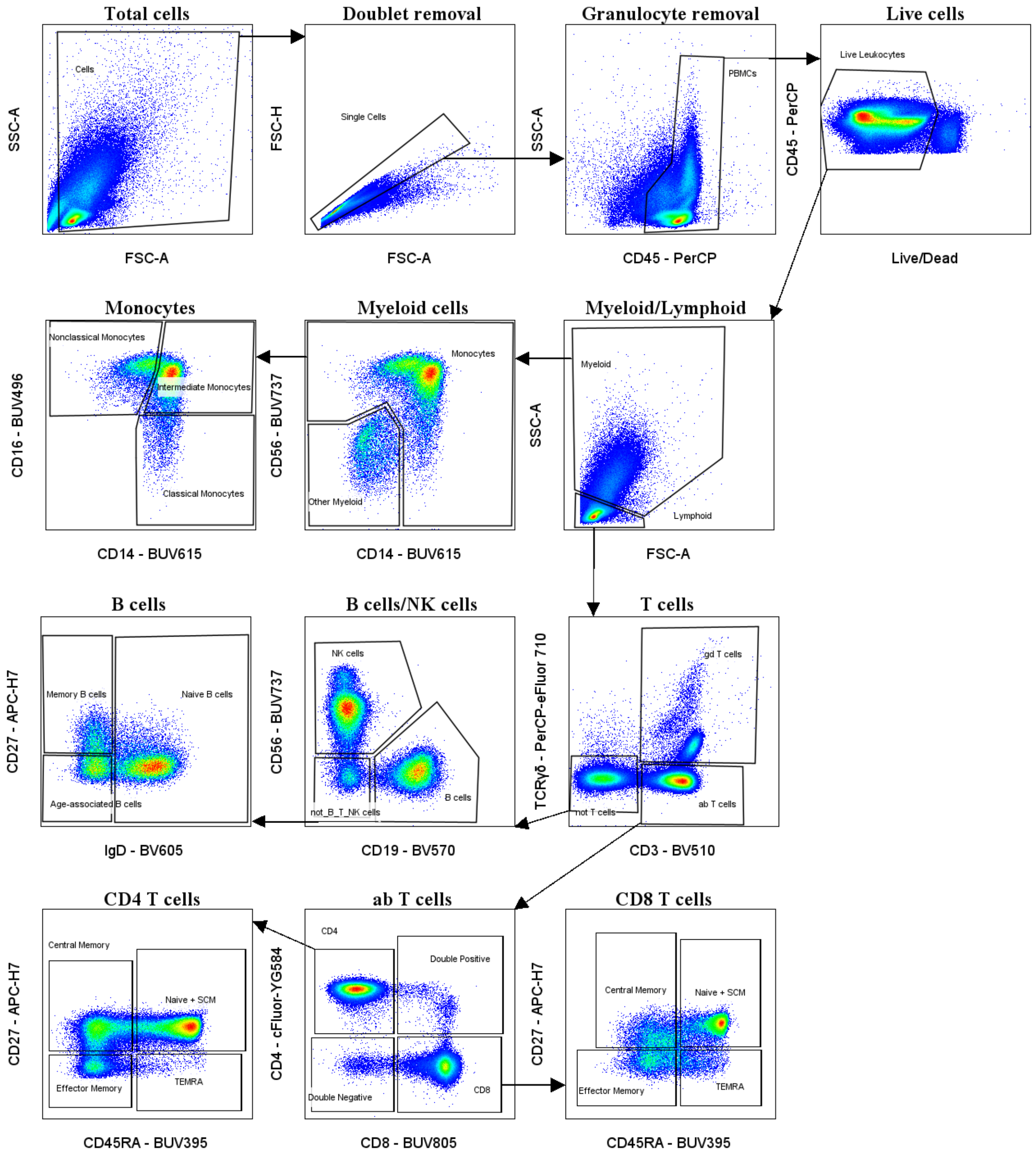


**Supplementary Fig. 7**: Gating strategy for flow cytometry immunophenotyping of PBMCs
NK, Natural killer, SCM, Stem cell memory; TEMRA, CD45RA+ T effector memory.

Supplementary Fig. 8 (related to Fig.4)

Metabolic parameters were calculated from the geometric mean fluorescence intensity (MFI) of the anti-puromycin antibody signal with each inhibitor relative to the control:

Co = Control treatment

DG = 2-Deoxy-D-Glucose treatment

O = Oligomycin A treatment

DGO = DG+O treatment

- $Glucose\ Dependence\ (\%) = \frac{100(Co-DG)}{(Co-DGO)}$

- $Mitochondrial\ Dependence\ (\%) = \frac{100(Co-O)}{(Co-DGO)}$

- $FAO\ and\ AAO\ Capacity\ (\%) = 100 - (\frac{100(Co-DG)}{(Co-DGO)})$

- $Glycolytic\ Capacity\ (\%) = 100 - (\frac{100(Co-O)}{(Co-DGO)})$

**Supplementary Fig. 8**: SCENITH dependencies and capacities equations
